# Osteomacs support osteoclast-mediated resorption and contribute to bone pathology in a postmenopausal osteoporosis mouse model

**DOI:** 10.1101/2021.02.05.429872

**Authors:** Lena Batoon, Susan M. Millard, Liza J. Raggatt, Andy C. Wu, Simranpreet Kaur, Lucas W.H. Sun, Kyle Williams, Cheyenne Sandrock, Pei Ying Ng, Michal Bartnikowski, Vaida Glatt, Nathan J. Pavlos, Allison R. Pettit

**Author notes:** **Corresponding Author:** Allison R Pettit. Mater Research Institute - The University of Queensland, Translational Research Institute, 37 Kent Street, Woolloongabba, QLD, 4102.

## Abstract

Osteal macrophages (osteomacs) support osteoblast function and promote bone anabolism, but their contribution to osteoporosis has not been explored. While mouse ovariectomy models have been repeatedly used, variation in strain, experimental design and assessment modalities, have contributed to no single model being confirmed as comprehensively replicating the full gamut of osteoporosis pathological manifestations. We validated an ovariectomy model in adult C3H/HeJ mice and demonstrated that it presents with human post-menopausal osteoporosis features, including reduced bone volume in axial and appendicular bone and bone loss in both trabecular and cortical bone including increased cortical porosity. Bone loss was associated with increased osteoclasts on trabecular and endocortical bone and decreased osteoblasts on trabecular bone. Importantly, this OVX model was characterised by delayed fracture healing. Using this validated model, we demonstrated that osteomacs are increased post-ovariectomy on both trabecular and endocortical bone. Dual F4/80 (pan-macrophage marker) and TRAP staining revealed osteomacs frequently located near TRAP^+^ osteoclasts and containing TRAP^+^ intracellular vesicles. Using an *in vivo* inducible macrophage depletion model that does not simultaneously deplete osteoclasts, we observed that osteomac loss was associated with elevated extracellular TRAP in bone marrow interstitium and increased serum TRAP. Using *in vitro* high-resolution confocal imaging of mixed osteoclast-macrophage cultures on bone substrate, we observed macrophages juxtaposed to osteoclast basolateral functional secretory domains scavenging degraded bone by-products. These data demonstrate a role for osteomacs in supporting osteoclastic bone resorption through phagocytosis and sequestration of resorption by-products. Finally, using *Siglec1* knockout mice, we demonstrated that loss of the macrophage-restricted molecule Siglec-1/CD169 is sufficient to cause age-associated low bone mass, emphasizing the macrophages, independent of osteoclasts, contribute to optimal skeletal health. Overall, our data expose a novel role for osteomacs in supporting osteoclast function and provide the first evidence of their involvement in osteoporosis pathogenesis.

## INTRODUCTION

Osteoporosis is a common and serious metabolic bone disease characterised by reduced bone volume, mineral density and quality ^(1)^. Fragility fracture is the clinical manifestation of osteoporosis and is associated with significant morbidity and increased mortality risk. About 30-50% of hip fracture patients are unable to live independently or walk without assistance ^(2)^. Furthermore, hip fractures have a one-year mortality rate of 20-33% ^(3,4)^ and this increased risk persists for up to 10 years ^(4)^. Despite advances in osteoporosis diagnosis and treatment options, fragility fracture remains a significant health challenge ^(5,6)^. This drives the need to improve understanding of the underlying pathophysiology of osteoporosis with the goal of achieving prevention or early intervention strategies.

Post-menopausal osteoporosis is the most common type of osteoporosis. Estrogen deficiency triggers an imbalance in bone remodelling, with bone resorption outpacing bone formation, leading to net bone loss. After menopause, the levels of biochemical markers of bone resorption are increased by 79-97% ^(7)^ while bone formation markers are either increased to a much lesser degree ^(7,8)^ or remain unchanged ^(9)^. Trabecular bone loss is significantly accelerated during perimenopause and post-menopause ^(10,11)^. In contrast, cortical bone remains relatively stable until after menopause, where cortical thinning and increased cortical porosity become evident^(10-13)^.

Ovariectomy (OVX) in rodents has been used extensively as a model of human post-menopausal osteoporosis. However, there remains no consensus regarding the optimal OVX model ^(14,15)^. In mice, OVX-induced skeletal impacts are influenced by genetic background, age at OVX and skeletal site analysed ^(16-21)^. While the C57Bl/6 strain is commonly employed due to the wide availability of genetically modified variants generated on this background, many of these studies performed OVX prior to 10 weeks of age at which time the skeleton is still actively growing ^(22)^. Consequently, these more accurately model estrogen impacts on attainment of peak bone mass ^(14)^. Notably, C57BL/6 mice have low peak bone mass compared to other strains ^(21)^ and females undergo spontaneous bone loss starting from 10-12 weeks of age ^(17,20,23)^, culminating in little residual trabecular bone for histomorphometric analysis at experimental end points ^(19,20)^. Therefore, both the choice of rodent strain and timing of OVX can have substantial impact on osteoporosis-relevant pathogenic mechanisms. While many animal studies have reported on specific hallmarks of osteoporosis, we were unable to identify any studies that comprehensively interrogated cellular changes, trabecular and cortical bone loss at multiple sites, biomarkers of bone turnover, strength properties and importantly pathology manifestations of osteoporosis. Improved confidence in the fidelity of mouse OVX models is needed through comprehensive investigation of whether they replicate the full spatiotemporal complexity of post-menopausal osteoporosis pathophysiology ^(14,24)^.

Estrogen-deficiency triggered macrophage dysfunction has been implicated in systemic immune activation contributing to the pro-inflammatory phenotype associated with osteoporosis and secretion of pro-osteoclastogenic cytokines ^(25-28)^. However, the potential role of osteal macrophages (osteomacs) in osteoporosis has not been explored. These bone resident tissue macrophages support osteoblast-mediated bone anabolism ^(29-37)^, providing a credible hypothesis that altered osteomac distribution and/or function could contribute to bone pathophysiology in osteoporosis.

The C3H/HeJ mouse strain is characterised by high and sustained peak bone mass ^(17,21,38)^ and exhibits robust trabecular bone loss at 4 weeks post-OVX ^(21)^. It is also the only strain reported to have a significant decrease in cortical area and width at 4 weeks post-OVX, and increased intracortical resorption spaces at 8 weeks post-OVX ^(18)^. Here, we performed extensive evaluation of OVX in the C3H/HeJ background to confirm that it is a robust model of post-menopausal osteoporosis, including investigation of whether it causes the critical pathological presentation of delayed fracture healing ^(14,39-42)^. Subsequently, we used this validated model of postmenopausal osteoporosis to assess the impact of estrogen deprivation on osteomac frequency and distribution that revealed a novel contribution of osteomacs in supporting osteoclast-mediated bone resorption.

## MATERIALS AND METHODS

### Experimental animals

Animal experiments were approved by The University of Queensland Health Sciences Ethics Committee and performed in accordance with the Australian Code of Practice for the Care and Use of Animals for Scientific Purposes. C57Bl/6 and C3H/HeJ mice were obtained from Australian Resources Centre (Canning Vale, WA, AU). Heterozygous *Siglec1*^tm1(HBEGF)Mtka^ (CD169^+/DTR^) knock-in mice ^(43)^ were sourced from Riken Bio Resource Centre (Yokohama, Kanagawa, Japan) and contain the human diphtheria toxin (DT) receptor (DTR) knocked into the murine *Siglec-1* locus. Homozygous *Siglec1*^tm1(HBEGF)Mtka^ (CD169-KO) mice were generated by crossing CD169^+/DTR^ mice. Male CD169-KO mice were utilized in this study. All mice were housed under specific pathogen-free conditions in the Translational Research Institute Biological Resource Facility, fed standard chow with *ad libitum* water access and simulated diurnal cycle. Dark Agouti rats were obtained from Australian Resources Centre and maintained in specific pathogen free facilities at The University of Queensland under protocols approved by the UQ Animal Ethics Unit.

### Ovariectomy and MouseScrew fracture surgery

Bilateral OVX was performed on skeletally mature 16-17-week-old female C3H/HeJ mice. Access to the ovaries was generated through a small incision in the abdominal wall from the bottom of the ribs to the top of the iliac crest. Ovaries were excised from the uterine horn followed by cauterisation to staunch bleeding and seal wound. Skin wound was closed using wound clips. For SHAM surgery, mice underwent the above procedure without ovary excision or cauterisation. A time course was undertaken with tissues collected at 1-, 2-, 3-, 4- or 8-weeks post-surgery. Fractures were generated in naïve 16-17-week-old female C3H/HeJ mice or C3H/HeJ mice 4 weeks post-OVX/SHAM surgery using the MouseScrew system as per manufacturer’s instruction (RISystems AG, CH). Specifically, mid-diaphyseal femoral fractures were created by blunt trauma and stabilised using an intramedullary screw. Tissues were collected at 1-, 3- or 5-weeks post-fracture surgery.

### *In vivo* osteomac depletion using CD169-DTR mice

Eight-week-old CD169-DTR male mice were randomly allocated to vehicle (0.9% sodium chloride) or DT (50µg/kg; MBL International Corporation, MA, USA) groups and treated via intra-peritoneal injection daily for 4 consecutive days. Tissues were collected 24 hours following the last treatment.

### *In vitro* macrophage and osteoclast assay and confocal immunofluorescence microscopy

M-CSF-dependent mouse bone marrow derived macrophages isolated from the long bones of C57BL/6 mice were cultured for 48 hours on devitalized bovine bone discs (0.2 mm x 6 mm) labelled with fluorescently conjugated inert bisphosphonate (Osteosense, Perkin Elmer, NEV10020EX) either in the absence or in the presence of mature osteoclasts that had been differentiated for 5-days under pro-osteoclastogenic conditions i.e. M-CSF (25 mg/ml, R&D) and RANKL (10ng/ml, R&D). Cells were then fixed with 4% paraformaldehyde and then stained with rat anti-Mouse F4/80, (BioRad Cl:A3-1) and goat anti-rat IgG cross-absorbed Alexa Fluor® 488 secondary antibody (ThermoFisher, A-11006) together with Rhodamine-conjugated Phalloidin (Invitrogen, R415) and Hoechst 33342 (Invitrogen, H3570) to visualize macrophages, filamentous (F)-actin and nuclei, respectively. Bone slices were mounted onto glass slides using ProLong™ Glass Antifade Mountant (ThermoFisher, P36980) and cells were imaged using a Nikon-A1 point scanning confocal microscope equipped with a PlanApo 100X oil objective lens (N.A.1.45). Images were collected using the systems NIS-C Elements software (Nikon) and processed using ImageJ software (NIH).

### Flow cytometry

Bone marrow from the right femora was collected for flow cytometry as previously described ^(30)^. Bone marrow cell suspensions were collected in 2% FCS in PBS and red and white blood cell differentials were analyzed using Mindray BC-5000 Auto Hematology Analyser (Vepalabs, QLD, AU). For flow cytometry analysis, cell suspensions were incubated with the appropriate antibody cocktail for 40 minutes in the dark, on ice with agitation. The osteoclast precursor antibody cocktail (all sourced from Biolegend) was made in Fc (fragment crystallizable antibody region) block and contained anti-B220-FITC (clone RA3-6B2), anti-CD3-FITC (clone 145-2C11), anti-Ter-119-biotin, anti-CD115-biotin (clone AFS98), anti-Ly6C-APCFire750 (clone HK1.4), and anti-CD11b-BV510 (clone M1/70). Cells were then washed and incubated with streptavidin-PE-BV785 for 30 minutes. Cells were washed and resuspended in 2% FCS in PBS for analysis. Specificity of staining was determined by comparison to unstained cells and appropriate isotype control cocktails. Five minutes prior to analysis 5 μg/ml 7-amino actinomycin D (Life Technologies, CA, USA) was added for exclusion of dead cells. Cells were analyzed using a CytoFLEX flow cytometer (Beckman Coulter, CA) and FlowJo version 10 (Tree Star Data Analysis Software).

### Immunohistochemistry (IHC), histological staining and histomorphometry methods

Mouse and rat bones were fixed in 4% paraformaldehyde (PFA) and processed for immunohistochemical analysis including decalification prior to processing as previously described ^(30,44)^. IHC on mouse bones was performed on deparaffinized and rehydrated sections with unconjugated primary antibodies against F4/80 (clone Cl:A3-1; Novus Biological, CO, USA), osteocalcin (polyclonal; Enzo Life Sciences, NY, USA), or relevant isotype control antibodies [normal rat IgG2b (Biolegend, CA, USA) and normal rabbit IgG (Santa Cruz Biotechnology, Inc., Santa Cruz, CA, USA) (Supplemental Figure 1) as previously described ^(30)^. IBA1 staining on rat bones was performed as per previously ^(44)^. All sections were counterstained with Mayer’s hematoxylin as per manufacturer’s instruction (Sigma-Aldrich, MO, USA). Tartrate-resistant acid phosphatase (TRAP) activity was detected as previously described ^(45)^. For dual staining for macrophage markers with TRAP, F4/80 or IBA1 staining was performed before TRAP detection. Staining was imaged using either VS120 slide scanner (Olympus, TYO, JP), and analyzed using Visiopharm® software (Visiopharm, DK) or Olympus BX50 microscope *via* Olympus CellSens standard software 7.1 (Olympus).

**Figure 1.**
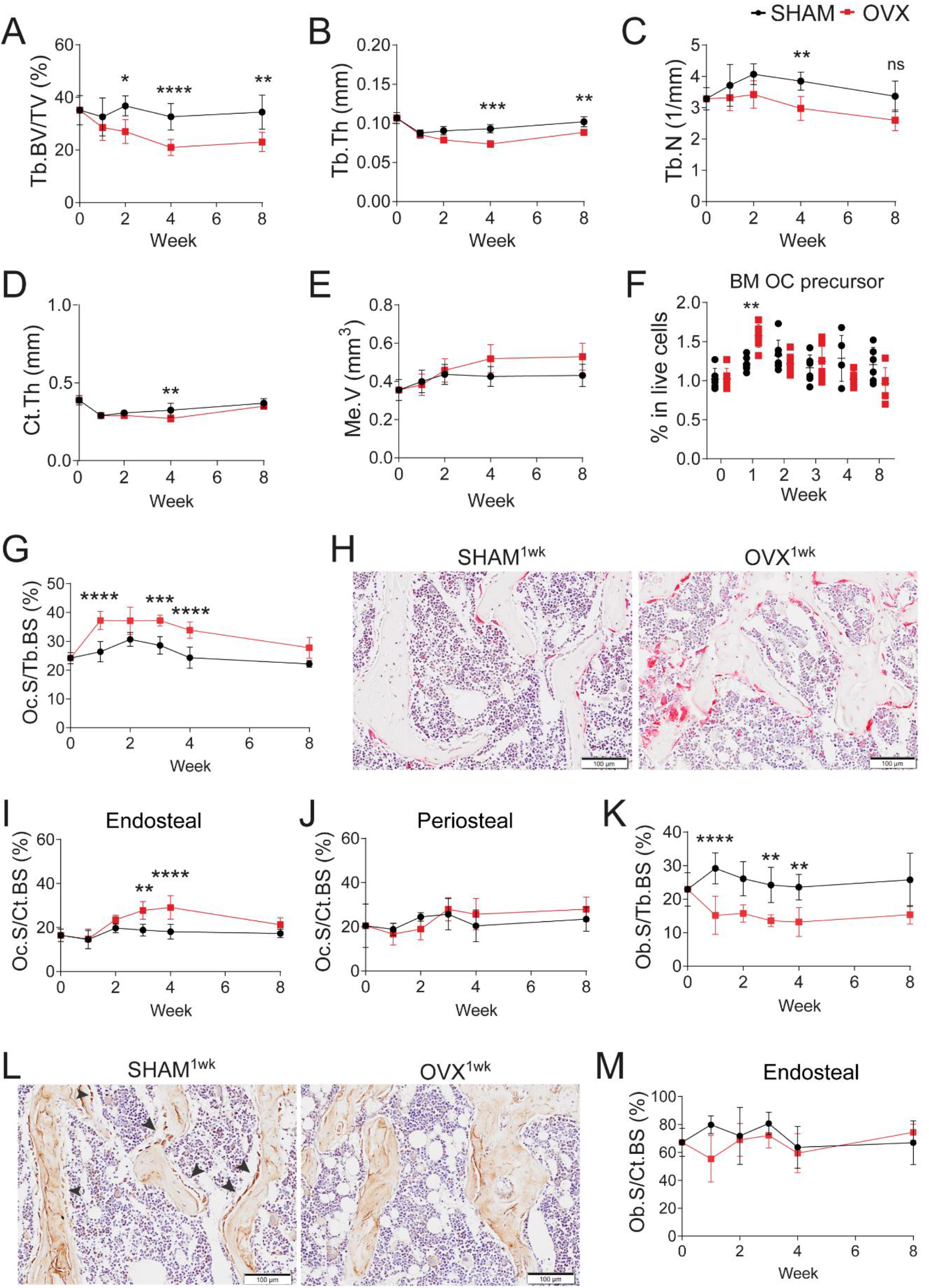
OVX in C3H/HeJ replicates features of post-menopausal osteoporosis. **A-C:** Micro-CT analyses of trabecular (Tb) bone volume (BV) represented as a ratio of total tissue volume (TV) (**A**); thickness (Th) (**B**); and number (N) (**C**) in distal femoral metaphysis of SHAM (black line) and OVX-operated mice (red line) over an 8-week time course post post-surgery. **D-E:** Micro-CT analyses of femoral mid-diaphysis showing changes in cortical (Ct) thickness (**D**) and medulla volume (Me.V) (**E**). Refer to Supplemental Figure 2 for regions of interest of trabecular and cortical bone compartments. **F:** Flow cytometry analysis of osteoclast (OC) precursors in the bone marrow (BM) represented as percent of live (7AAD^neg^) cells and gated as CD3^neg^Ter119^neg^B220^neg^CD115^+^CD11b^neg^Ly6C^+^ cells. **G:** Histomorphometry quantification of percent TRAP^+^ osteoclast surface (Oc.S) per trabecular bone surface (Tb.BS). **H:** Representative images of TRAP staining (magenta) in sagittal sections of trabecular bone in SHAM and OVX-operated animal at 1-week post-surgery. **I-J:** Histomorphometry quantification of percent length of TRAP^+^ osteoclast surface (Oc.S) per cortical bone surface (Ct.BS) on endosteal (**I**) and periosteal (**J**) surfaces. **K:** Histomorphometry quantification of percent osteocalcin^+^ osteoblast surface (Ob.S) per trabecular bone surface (Tb.BS). **L:** Representative images of osteocalcin staining showing osteoblasts (dark brown; arrows) on trabecular bone in SHAM and OVX-operated mice at 1-week post-surgery. **M:** Histomorphometry quantification of percent length of osteocalcin^+^ osteoblast surface (Ob.S) per endocortical bone surface (Ct.BS). In **G, I, J, K, M**, each data point represents the mean of 3 independent measurements at 3 different sectional depths at least 50 μm apart. Statistical significance was determined using two-way ANOVA with Tukey’s post-test. ****, p < 0.0001; ***, p < 0.001; **, p < 0.01; *, p < 0.05. Error bars represent standard deviation. Original magnification: 200X; Scale bar: 100 µm.

The length of F4/80^+^ osteomac (F4/80^+^ cells within 3 cell diameter of bone) surface per total bone surface, osteocalcin^+^ osteoblast surface per total bone surface and TRAP^+^ osteoclast surface per total bone surface were quantified on endocortical and trabecular bone in a blinded manner. On cortical bone, the measurements were taken between 1.5 mm from the base of the growth plate and 4.5 mm distally from this point. A total length of 3.3 ± 0.5 mm of endocortical bone surface was examined. On trabecular bone, the average total bone surface examined for all parameter was 8.9 mm ± 2.4 mm. The number of osteocyte nuclei and TRAP^+^ osteocytes in cortical bone were enumerated using an automated approach using Visiopharm software. The generated trained algorithm detected nuclei (blue), TRAP staining (magenta) and ‘other’ unstained areas which include the bone and blood vessel. A total area of 1.7 mm^2^ ± 0.3 mm^2^ of cortical bone was examined per section.

### Micro-CT imaging and reconstruction

Bones assessed by micro-CT were transferred to phosphate-buffered saline (PBS) following fixation in 4% PFA without undergoing any decalcification process. Eight-week-old CD169-KO bones were scanned using a desktop µCT imaging system (µCT40; Scanco Medical, CH) while the other bones were scanned using Bruker’s Skyscan 1272 (Bruker microCT, BE). The x-ray settings were standardised to 70 kV and 142 µA. The x-ray filter used was a 0.5 mm aluminium. The entire femur and second lumbar vertebrae were scanned over 360° rotation in 0.8° rotational steps and the exposure time was set to 470 ms. Projections were acquired with nominal resolutions of 10 µm and each slice contained 1224 x 820 pixels. For intracortical porosity assessment, high-resolution imaging was employed. The mid-diaphyseal region of the femur was scanned at a rotation step of 0.25° over 360° rotation and with an exposure time of 1400 ms. The projections acquired were at 2.7 µm nominal resolution with each slice containing 2452 x 1620 pixels.

All X-ray projections were reconstructed using a modified back-projection reconstruction algorithm (NRecon 1.7.3.1 software-SkyScan, Bruker) to create cross-sectional images and visualized in 3D using CTvox 3.3.0 (Bruker). Reconstruction parameters included ring artefact correction (2-6), beam hardening correction (40-50%), and misalignment correction. Reconstruction was performed in a blinded manner. Reconstructed images were then analysed using CTAn 1.19 software (Bruker) which has inherent 2D and 3D analysis tools. Regions of interest examined in the trabecular and cortical bone compartments are exemplified in Supplemental Figure 1C. For MouseScrew fractures, the total callus formed was examined through automated greyscale intensity-based algorithm that distinguished new bone formed (hard callus) from original cortical bone. For intracortical porosity, analysis was performed as per previously (Hemmatian et al., 2017) with slight modification. Intracortical porosity was segmented based on size to delineate between noise, osteocyte lacunae and intra-cortical canals (nerves and blood vessels). Objects less than 100 µm^3^ were considered to be noise, objects between 100-1400 µm^3^ were assigned as osteocyte lacunae and objects greater than 1400 µm^3^ were assigned as intra-cortical canals. These volume limits were based on previous studies that assessed cortical porosity using high resolution micro-CT ^(46,47)^.

### Fracture cortical bridging score

Blinded assessment of fracture cortical bridging was performed by manual examination of 2D micro-CT images generated using DataViewer (Bruker). Fracture sites were assessed at 2 randomly selected transverse slices. At each of these transverse planes, 5 different depths (100 µm apart) were assessed in both the coronal and sagittal plane achieving comprehensive fracture site sampling for scoring of fracture gap bridging. Semi-quantitative scores were as follows: 2, both cortices were bridged; 1, one cortex was bridged; and 0, neither cortex was bridged.

## Statistical analyses

Statistically significant differences were determined using either a two-way ANOVA with Tukey’s multiple comparison post-test or two-tailed unpaired t-test in PRISM 8 (GraphPad Software, CA, USA). Group sizes are indicated in graphs and/or figure legends. A value of p < 0.05 was considered statistically significant.

## RESULTS

### OVX in adult C3H/HeJ mice caused osteopenia that replicated human osteoporosis pathophysiology

While OVX in the C3H/HeJ strain has been used as a mouse model of post-menopausal osteoporosis ^(17,18,21,48)^, the full spectrum of pathophysiological manifestations have not been comprehensively examined in this, or any other rodent strain, within a single study to confirm fidelity as a robust model of osteoporosis ^(20)^. As experimental design (i.e. age at OVX and experimental endpoints) often vary between studies, this creates ambiguity and variation in the literature ^(14,20)^. OVX-induced bone pathology in the C3H/HeJ background was characterised over an 8-week time course post-OVX in skeletally mature 16-17-week-old mice. Uterine atrophy was evident as early as 1-week post-OVX (Supplemental Figure 2A). Bone marrow adiposity was elevated at 4-weeks post-OVX (Supplemental Figure 2B), which is a known feature of human post-menopausal osteoporosis ^(49,50)^. Bone volume was assessed in femoral distal metaphysis and mid-diaphyseal shaft (Supplemental Figure 2C). OVX resulted in reduced metaphyseal trabecular bone volume fraction as early as 2-weeks post-surgery and this was maintained up to 8 weeks (Figure 1A and Supplemental Figure 2D). Trabecular bone thickness and number were also reduced at 4-weeks post-OVX (Figure 1B-C). Diaphyseal cortical bone volume was unchanged (Supplemental Figure 2E) but cortical thickness was significantly reduced at 4-weeks post-surgery (Figure 1D), and there was a trend toward increased medullary volume (Figure 1E). Significant bone loss was also evident in vertebrae from 1-week post-surgery (Supplemental Figure 2F-H). Although serum C-telopeptide of type I collagen (CTX-1) and TRAP were not different between SHAM- and OVX-operated mice (Supplemental Figure 2I-J), monocyte chemokine attractant protein 1, which contributes to elevated osteoclastogenesis ^(51)^ and has been shown to be increased in osteoporotic patients ^(52)^, was increased post-OVX (Supplemental Figure 2K).

**Figure 2.**
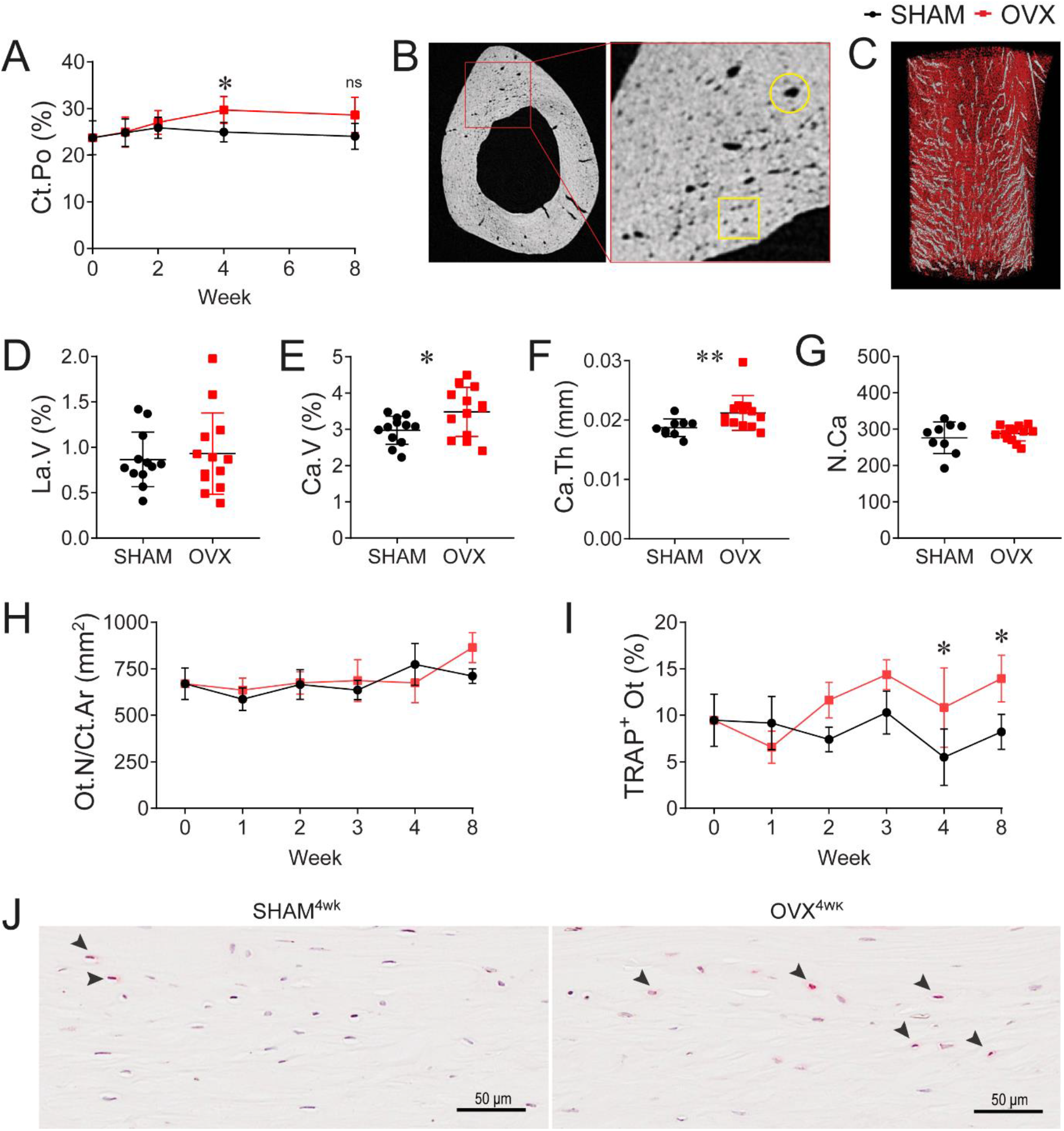
OVX increased cortical porosity and TRAP+ osteocytes. **A:** Micro-CT analyses of femoral mid-diaphysis to determine cortical porosity at 10 µm resolution across an 8-week time course post-SHAM or OVX surgery. **B:** Micro-CT 2D slice imaging at 2.7 µm resolution at 4 weeks post-surgery exemplifying size-based approach to distinguish osteocyte lacunae (yellow square) and vascular/nerve canals (yellow circle). **C:** Micro-CT 3D reconstruction showing segmentation based on size with osteocyte lacunae in red and canals in grey. **D-G:** Quantification of lacunae volume (La.V) (**D**), canal volume (Ca.V) (**E**), canal thickness (Ca.Th) (**F**) and canal number (N.Ca) (**G**) at 2.7 µm resolution micro-CT imaging at 4 weeks post-surgery. **H-I:** Histomorphometry quantification of total osteocyte number per cortical area (Ot.N/Ct.Ar) (**H**) and frequency of TRAP^+^ osteocytes (%TRAP^+^ Ot) (**I**) in cortical bone. For **H** and **I**, each data point represents the mean of 2 independent measurements at 2 different sectional depths at least 50 μm apart. **J:** Representative images of TRAP-stained sections of bone from SHAM and OVX-operated mice at 4 weeks post-surgery showing TRAP^+^ osteocytes (arrowheads). Statistical significance was determined using two-way ANOVA with Tukey’s post-test (**A, H, I**) or two-tailed unpaired t-test (**D-G**). *, p < 0.05; **, p < 0.01. Error bars represent standard deviation. Original magnification: 400X; Scale bar: 50 µm.

A transient but significant increase in bone marrow osteoclast precursor cells was detected at 1-week post-OVX (Figure 1F). Osteoclast surface per trabecular bone surface was elevated from 1-through 4-weeks post-OVX but returned to equivalent levels with SHAM-operated mice at 8 weeks (Figure 1G-H). TRAP^+^ osteoclasts were also increased on the endocortical surface at 3 and 4 weeks and normalised at 8 weeks post-OVX (Figure 1I). TRAP^+^ osteoclast number on the periosteal surface remained unaltered during the time course (Figure 1J). Conversely, osteocalcin^+^ osteoblast number on trabecular bone was significantly reduced over the first 4-weeks post-OVX (Figure 1K-L). Osteoblast frequency on the endocortical surface was unaffected (Figure 1M).

Increased intracortical porosity is an important determinant of bone fragility that underlies fracture risk in osteoporosis ^(12)^. Increased intracortical porosity was evident 4-weeks post-OVX (Figure 2A) and higher resolution imaging demonstrate that this was not due to changes in osteocyte lacunar volume (Figure 2E) but increased total intra-cortical canal volume (Figure 2F). There was no change in intra-cortical canal number, but thickness was increased (Figure 2F-G). Analysis of TRAP-stained sections from OVX- and SHAM-operated mouse bones showed no difference in osteocyte number (Figure 2H) but a significant increase in TRAP-expressing osteocytes (Figure 2I-J). Overall, these results confirm that OVX in C3H/HeJ recapitulates the major cellular adaptations and bone volumetric changes characteristic of a high remodelling turnover osteoporosis.

### Fracture in ovariectomised C3H/HeJ mice is associated with delayed bone healing

While spontaneous fracture has not been observed in OVX rodent models ^(15)^, an important pathological manifestation of osteoporosis is delayed fracture healing ^(14)^. Modelling of this has proven challenging in small animals, particularly in mice, but this has not been previously examined after OVX in C3H/HeJ mice. Fracture fixation quality and reproducibility are a paramount consideration in osteopenic bone, and these were addressed by using the MouseScrew mid-diaphyseal fracture model. The fracture generated in this model is bridged by 5-weeks in naive adult C3H/HeJ female mice with healing progressing via periosteal endochondral callus formation (Figure 3A). At 1-week, the callus was predominantly cartilage and granulation tissue, while at 3-weeks the callus was largely bone (Figure 3A-B). At 5-weeks, the callus had undergone extensive remodelling to woven bone that contained pseudo-bone marrow (Figure 3A-B).

**Figure 3.**
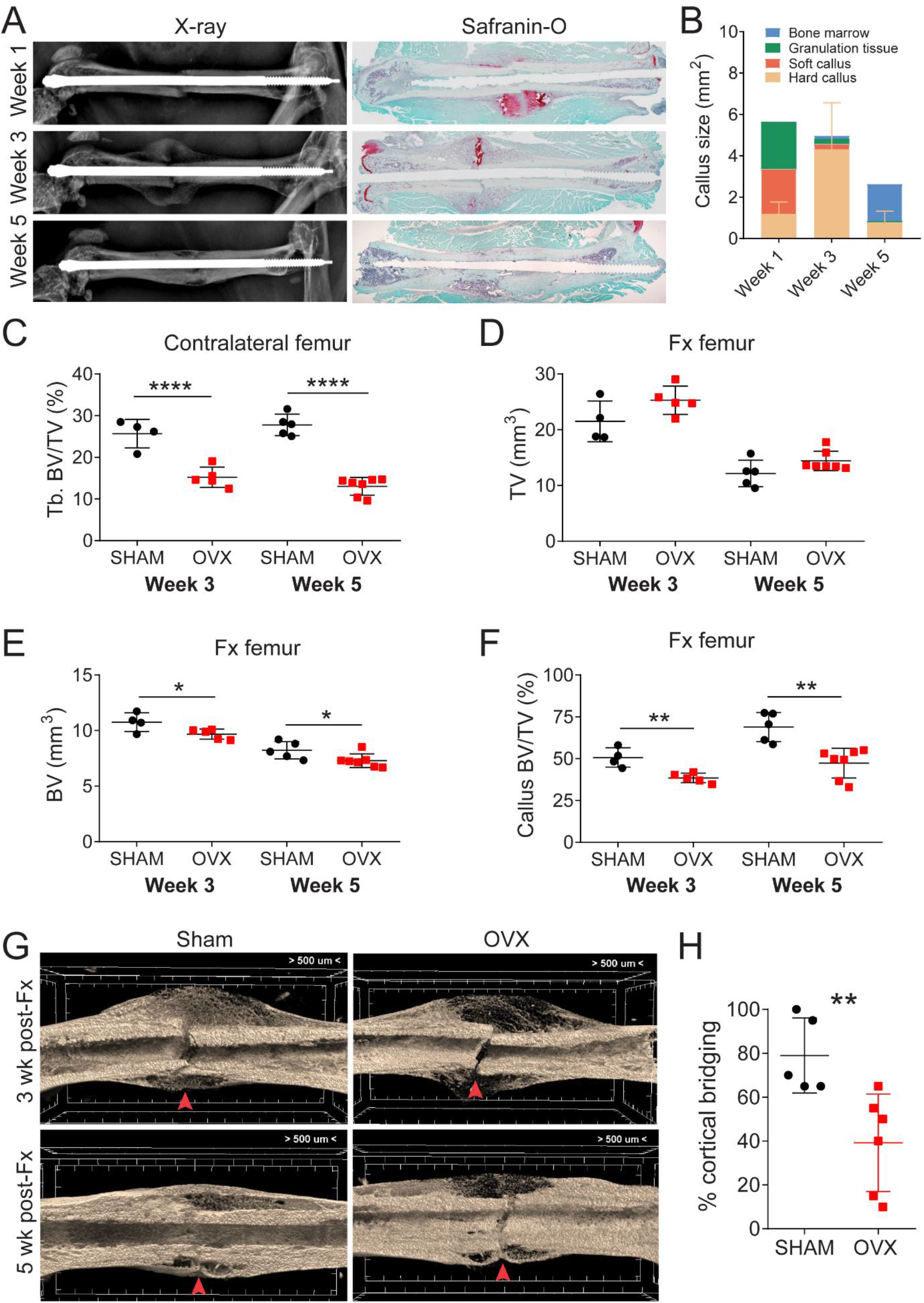
Delayed fracture healing in ovariectomised C3H/HeJ mice. **A:** Representative x-ray images (left) and safranin-O staining (right) of fractured femur from naïve mice at 1-, 3- and 5-weeks after MouseScrew fracture. Cartilage tissues stain red. Original magnification: 40x. **B:** Fracture composition at 1-, 3- and 5-weeks post-fracture showing amount of bone marrow, granulation tissue, soft callus and hard callus formed at different experimental endpoints in naïve mice. **C:** Micro-CT analysis of the contralateral unfractured femur of SHAM and OVX-operated mice showing percent trabecular (Tb) bone volume per total volume (BV/TV) at 3- or 5-weeks post-fracture. **D-F:** Micro-CT analysis of the fracture callus showing total callus volume (TV, tissue volume) (**D**), bone volume (BV) within the callus (**E**) and percent bone volume per total callus volume (BV/TV) (**F**) at 3- or 5-weeks post-fracture in SHAM- or OVX-operated mice. **G:** Representative 3D reconstructions of the fracture sites (red arrowheads) and associated callus in SHAM- and OVX-operated mice at 3- or 5- weeks post-fracture. **H:** Percent cortical bridging of the fracture gap at 5 weeks post-fracture. Statistical significance was determined using two-tailed unpaired t-test comparing OVX-with SHAM - operated mice at each experimental endpoint. *, p < 0.05; **, p < 0.01, ****, p < 0.0001. Error bars represent standard deviation.

Based on our characterisation of the OVX-induced osteopenic phenotype (Figure 1A-E) MouseScrew fractures were generated at 4 weeks post-OVX. OVX-associated trabecular bone loss was confirmed in the unfractured contralateral femur at the experimental endpoint (Figure 3C). Tissues were harvested at 3- and 5-weeks post-fracture to coincide with peak bone formation and cortical union, respectively (Figure 3A-B). Although total callus volume was unchanged between groups (Figure 3D), the bone volume fraction within the callus was significantly reduced in OVX-operated mice at both post-fracture time points (Figure 3E-G). Importantly, the cortical fracture gap at 5-weeks post-fracture revealed significantly reduced bridging in OVX mice compared to the SHAM controls (Figure 3G-H). These results confirmed delayed bone repair associated with OVX in C3H/HeJ mice. Replication of the key pathological complication of delayed fracture repair validates OVX in C3H/HeJ mice as a high-fidelity model of osteoporosis.

### Osteomacs are increased post-OVX and contribute to clearance of resorption by-products

To extend understanding of OVX pathophysiology, we assessed the impact of OVX in C3H/HeJ mice on osteomac homeostasis. OVX increased F4/80^+^ osteomac surface per bone surface on both trabecular and endocortical bone surfaces (Figure 4A-D). Osteomacs were increased on trabecular bone surfaces from 2-weeks post-OVX and remained elevated across the full-time course (Figure 4A) but were only transiently increased at 4-weeks on the endocortical bone surface. This paralleled osteoclast dynamics at these anatomical locations (Figure 1G and I). In both SHAM- and OVX-operated mice, F4/80^+^ osteomacs were often positioned near, or directly adjacent to, the basolateral membrane of TRAP^+^ osteoclasts on the trabecular bone surface (Figure 4E). The osteoclast-basolateral membrane contains the functional secretory domain (FSD) and so it was of interest that TRAP^+^ intracellular vesicles were evident inside osteoclast-adjacent osteomacs (Figure 4E). To determine whether this phenomenon occurred in another species, we developed a protocol that independently identified osteoclasts and osteomacs in rat bones via dual TRAP and IBA1 staining. Similar to observations in mouse, IBA1^+^ osteomacs were present in close proximity to TRAP^+^ osteoclasts in naïve growing bone at sites of remodelling. Again, these osteoclast-adjacent osteomacs also contained TRAP^+^ intracellular vesicles (Figure 4F).

**Figure 4.**
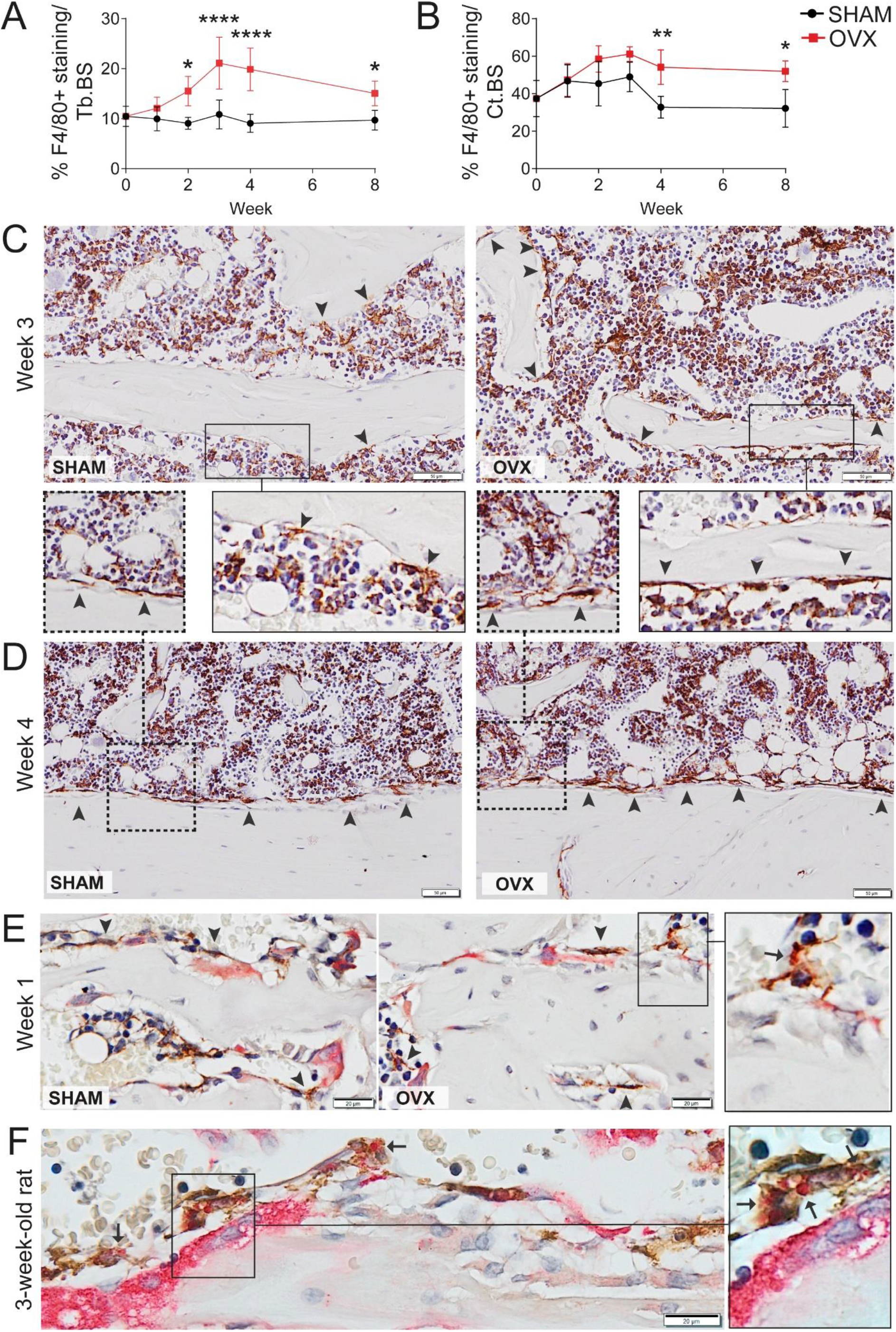
Osteomacs increased post-OVX in both trabecular and cortical bone. **A-B:** Histomorphometry quantification of F4/80^+^ osteomacs on trabecular (Tb) (**A**) and endocortical (Ct) (**B**) bone surfaces (BS). Each data point represents the mean of 3 independent measurements at 3 different sectional depths at least 50 μm apart. Statistical significance was determined using two-way ANOVA with Tukey’s post-test. ****, p < 0.0001; **, p < 0.01; *, p < 0.05. Error bars represent standard deviation. **C-D:** Representative images of sections stained with F4/80 in the trabecular bone region at 3 weeks post-surgery (**C**) and endocortical bone region (**D**) at 4 weeks post-surgery of SHAM or OVX-operated mice. Osteomacs on bone surfaces are indicated by arrowheads. **E:** Representative images of dual staining for F4/80 and TRAP activity in SHAM or OVX-operated mice at 1-week post-surgery showing F4/80^+^ osteomacs located adjacent to TRAP^+^ osteoclast (left and middle panel, arrowheads) and intracellular TRAP^+^ vesicles within a F4/80^+^TRAP^neg^ osteomac (right panel, arrows). **F:** Representative image of dual staining for the rat macrophage marker IBA1 and TRAP activity on 3-week-old rat tibia showing IBA1 osteomacs adjacent to osteoclasts containing distinct intracellular TRAP^+^ vesicles within their cytoplasm (arrows). Original magnification: 400X. Scale bar: 50 µm (**C-D**) and 20 µm (**E-F**).

Based on these in situ observations, we investigated whether osteomacs are involved in clearance of excess TRAP expelled via the FSD during the process of osteoclast-mediated bone resorption. The CD169-DTR mouse model was used to deplete osteomacs/macrophages without off-target impacts on osteoclasts ^(30)^. Of note, we have previously shown using this model that depletion of osteomacs/macrophages resulted in significantly increased serum TRAP without accompanying changes in osteoclast number, area or function ^(30)^. A 4-day DT-treatment regimen achieved profound loss of bone marrow macrophages as well as osteomacs on both endocortical (Figure 5B) and trabecular (Figure 5C and E) bone surfaces. Osteoclasts on the trabecular bone were unaffected (Figure 5D and F). In vehicle control, intracellular TRAP^+^ vesicles were observed in TRAP negative cells that were close to the bone surface (Figure 5D, arrowheads). Whereas in DT-treated mice, there was an increase in anuclear extracellular TRAP^+^ deposits within the bone marrow (Figure 5G and D). Together these findings indicate that, in the absence of CD169^+^ osteomacs/macrophages, excess TRAP accumulates in bone marrow extracellular fluid and perfuses into the circulation.

**Figure 5.**
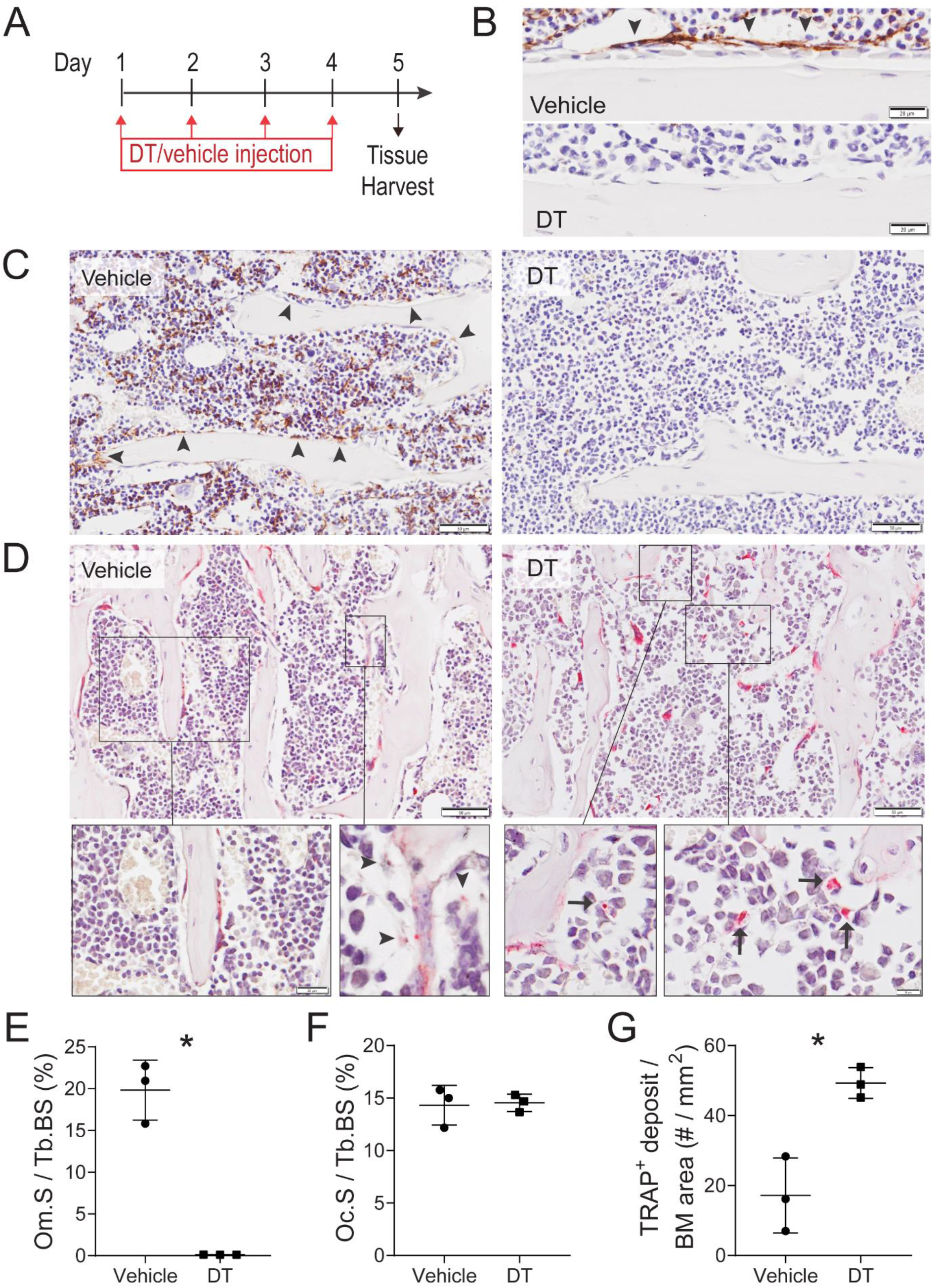
CD169+ osteomac/macrophage depletion increased TRAP+ deposits within bone marrow interstitium. **A:** Schematic of experimental design showing DT/vehicle injection regimen and tissue harvest. **B-C:** Representative images of IHC staining for F4/80 showing osteomacs abundant in the vehicle-treated control but almost absent in DT-treated mice on both endocortical (**B**, arrowheads) and trabecular bone surfaces (**C**, arrowheads). **D:** Representative images of TRAP staining (magenta) in metaphyseal bone region in vehicle or DT-treated mice showing TRAP subcellular vesicle within TRAP^neg^ cells in vehicle-treated mice (arrowhead) and interstitial TRAP^+^ deposit in DT-treated mice (arrow). **E:** Histomorphometry quantification of F4/80^+^ osteomacs surface (Om.S) per trabecular bone surface (Tb.S). **F-G:** Histomorphometry quantification of osteoclast surface (Oc.S) per trabecular bone surface (Tb.S, **F**) and TRAP^+^ deposit in the bone marrow (BM, **G**) in vehicle and DT-treated mice. Each data point represents the mean of 3 independent measurements at 3 different sectional depths at least 50 μm apart. Statistical significance was determined using two-tailed unpaired t-test. *, p < 0.05. Error bars represent standard deviation. Original magnification: 400X. Scale bar: 20 µm (**B** and **D** bottom left panel), 50 µm (**C** and **D** top panels), 10 µm (**D** bottom right panel).

In accordance with these *in vivo* observations, *in vitro* coculture of macrophages and osteoclasts on fluorescently labelled devitalised bone surfaces demonstrated direct interaction between macrophages and multinucleated bone-resorbing osteoclasts (Figure 6A). In particular, we frequently observed direct membrane contacts and projections between macrophages and the basolateral-orientated functional secretory domain (FSD) of osteoclasts, i.e. the site where degraded bone particulate (cyan colour) is expelled ^(53)^ (Figure 6A). Curiously, we noted that macrophages juxtaposed to the osteoclast FSDs also contained fluorescently labelled bone particles (cyan colour) within their cytosol suggestive of ‘opportunistic’ phagocytosis of resorption by-products by nearby macrophages. Indeed, this bone particulate uptake was only observed when macrophages were cocultured directly with bone-resorbing osteoclasts (Figure 6A) but not when macrophages where cultured on bone surfaces alone (Figure 6B). This implies that the bone particles within macrophages in co-cultures are not a result of direct resorption by macrophages but rather reflect the indirect uptake of resorption by-products liberated by osteoclasts.

**Figure 6.**
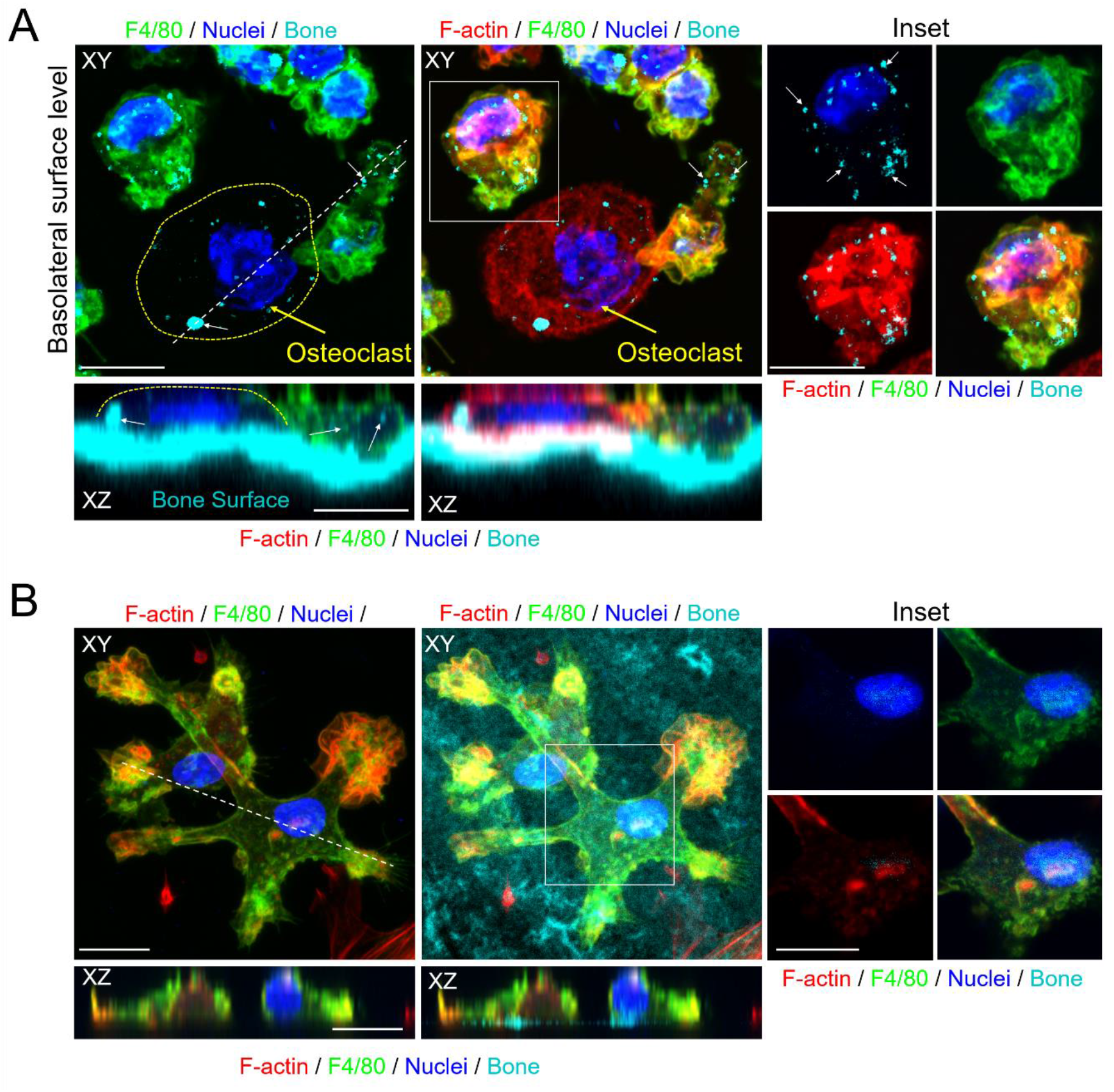
Macrophages accumulate resorption by-products liberated from bone by osteoclasts. **A-B:** Fluorescent confocal images of a mouse osteoclast and macrophage co-culture (**A**) and mouse macrophage monoculture (**B**) grown on devitalized bovine bone discs fluorescently labelled with osteosense (cyan). Red represents F-actin (Rhodamine-conjugated Phalloidin), green represents F4/80 expression and blue represents nuclei (Hoechst 33342). XY?=?top view/osteoclast basolateral functional secretory domain (top left and middle panels), XZ?=?side view (bottom panels). Insets are magnified images of the macrophage in boxed region. ‘Circular’ dotted line outlines osteoclast (yellow arrow) cellular boundary. White arrows denote internalized degraded bone particulate. Straight dashed line indicates the orientation of XZ plane shown in XY plane. Scale bar?= 10 µm.

### CD169-KO is associated with a low bone mass phenotype

Finally, we utilised CD169-KO mice to examine whether loss of this tissue resident macrophage-restricted molecule, that is expressed by osteomacs ^(30)^, impacted bone homeostasis. Frequency of blood and bone marrow suspension common myeloid cell population were minimally impacted by CD169 knockout (data not shown). Bone phenotype was examined in male mice CD169-KO to avoid confounding impacts of spontaneous bone loss in female mice on this background ^(17,20,23)^. In 8-week-old male CD169-KO mice, we observed a significant but transient reduction in osteomacs on trabecular bone surfaces (Figure 7A). There was a trend toward decreased osteoblast frequency in CD169-KO male mice at 8 weeks and this decline reached statistical significance at 16 weeks (Figure 7B-C). Osteoclast surface per trabecular bone surface was not different between groups at 8 weeks but significantly increased by 16 weeks (Figure 7D-E). Reductions in trabecular bone volume and number (Figure 7F-G) with increased trabecular spacing (Figure 7H) were evident at 16 weeks indicative of age-associated low bone mass in CD169-KO male mice.

**Figure 7.**
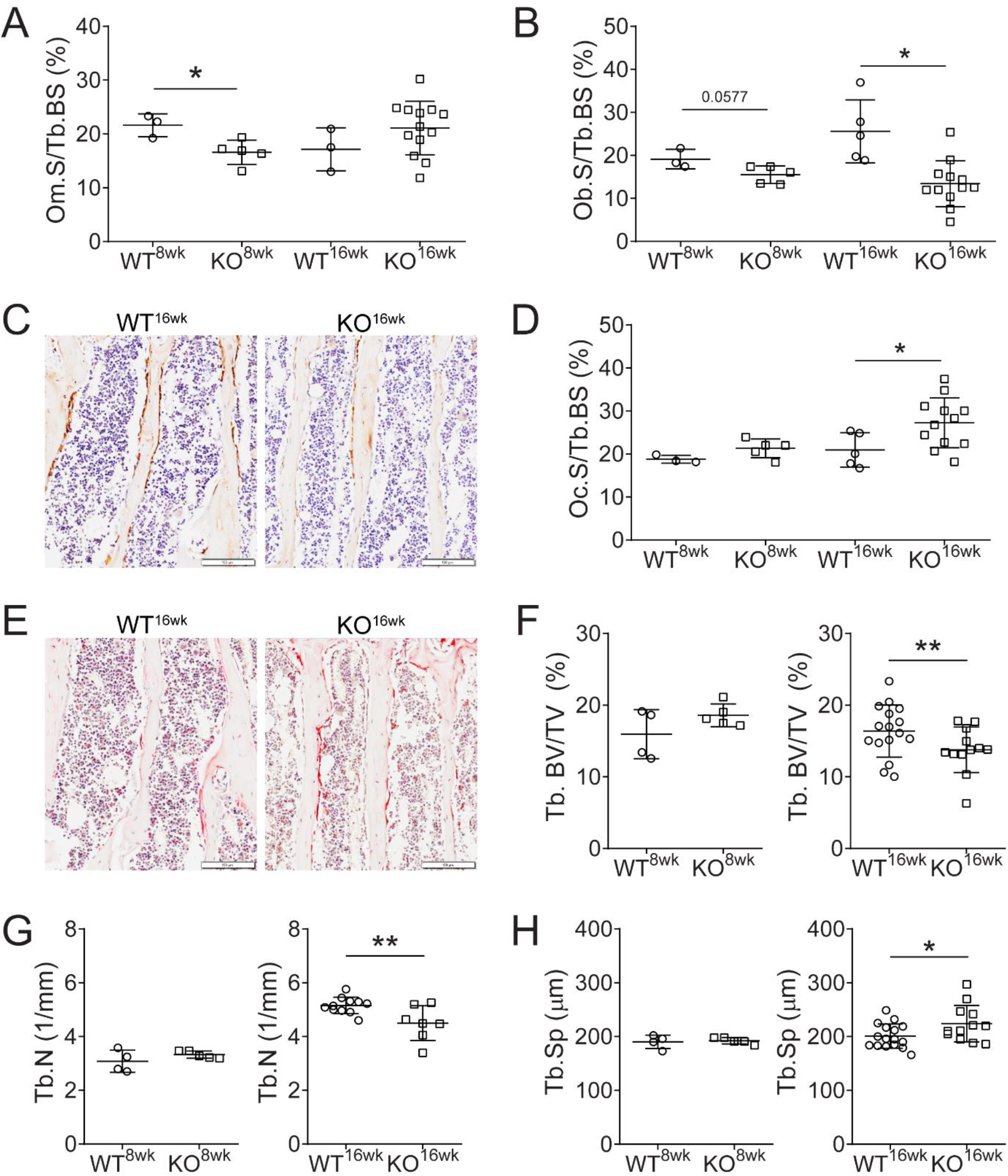
Deletion of macrophage-specific molecule CD169 results in low bone mass phenotype. **A:** Histomorphometry analysis of percent osteomac surface (Om.S) per trabecular bone surface (Tb.BS) in WT and CD169 KO mice at 8 and 16 weeks. **B-E:** Quantification of osteoblast surface (Ob.S) per trabecular bone surface (Tb.BS) (**B**) and osteoclast surface (Oc.S) per trabecular bone surface (Tb.BS) (**D**). Representatives images showing osteocalcin (**C**) and TRAP staining (**E**) on the trabecular bone. **F-H:** Micro-CT analysis of trabecular (Tb) bone volume fraction (BV/TV) (**F**), trabecular number (Tb.N) (**G**) and trabecular spacing (Tb.S) (**H**) in wild-type (WT) and CD169 KO mice at 8 and 16 weeks. Statistical significance was determined using two-tailed unpaired t-test comparing knockout mice with wild-type control at different ages. *, p < 0.05; **, p < 0.01. Error bars represent standard deviation. Original magnification: 400X. Scale bar: 50 µm.

## DISCUSSION

Osteomacs are key regulators of osteoblast-mediated bone formation in skeletal growth, homeostasis and regeneration, therefore, we investigated whether these resident tissues macrophages are impacted by estrogen deficiency and potentially contribute to osteoporosis pathophysiology. To explore this novel pathobiology and improve the translatability of the generated knowledge, we first undertook comprehensive validation of OVX in adult C3H/HeJ as a mouse model of post-menopausal osteoporosis. This OVX model replicated key features of osteoporosis including: decreased volume in axial and appendicular bones; bone loss in both metaphyseal trabecular and diaphyseal cortical bone, including increased cortical porosity; elevated serum marker of osteoclastogenesis and increased osteoclast precursors in the periphery; and, reduced trabecular osteoblasts and increased osteoclasts in both trabecular and cortical bone. Importantly, we also demonstrated that this OVX model was characterised by delayed fracture healing, which is an important complication of post-menopausal osteoporosis fragility fractures ^(14)^. Osteomacs were increased in both cortical and trabecular bone and were frequently adjacent to the basolateral membrane of osteoclasts at sites of bone reportion. We subsequently provided evidence that osteomacs support osteoclast-mediated bone resorption via clearance of resorption by-products including bone particulate and TRAP. Finally, we demonstrated that loss of the macrophage-restricted molecule CD169 is sufficient to cause age-associated low bone mass phenotype, emphasizing the importance of macrophages in attaining or sustaining peak bone health.

We chose the C3H/HeJ strain in which to model osteoporosis as it attains and sustains a high peak bone mass ^(17,21,38)^ with previous studies providing confidence that both trabecular ^(21)^ and cortical bone ^(18)^ loss occur in this strain after OVX in adult mice. This is also the only mouse strain in which OVX has been reported to increase intracortical porosity ^(18)^. These are important osteoporosis hallmarks that should be replicated in a mouse model ^(12,14,54)^. We confirmed that both trabecular (distal femur and lumbar vertebra) and cortical bone volume is reduced after OVX in C3H/HeJ mice. Importantly, kinetics of bone loss in these regions paralleled human disease patterns with trabecular loss occurring early and reductions in cortical bone occurring later. Intracortical porosity was also observed herein which is an important feature of human osteoporosis as it impacts mechanical properties of bone ^(55)^. Intracortical porosity was due to enlargement not number of intracortical canals, likely through widening of existing blood vessel and nerve canals. This is similar to observations in humans ^(56)^, but differs from previous observation in a C57Bl/6 model of senile osteoporosis which suggested porosity was due to creation of new osteons-like structures ^(57)^. We observed increased number of TRAP^+^ osteocytes, a phenomenon that has previously been reported in an OVX-induced model of osteoporosis in rats ^(58)^. TRAP expression has been associated with osteocytic osteolysis in the context of lactation ^(59)^ although we did not observe an increase in lacunae size herein.

Further investigation is required to determine the osteocyte functional implications of TRAP expression after OVX in C3H/HeJ mice. We also confirmed that OVX in adult C3H/HeJ induced expected high bone turnover osteoporosis-associated alterations in osteoclast and osteoblast frequency and elevation of a serum marker of increased osteoclastogenesis. A limitation of this study was that we did not directly assess bone mechanical properties after OVX, although reduced bone strength and stiffness was reported by Li et. al. in C3H/HeJ mice using a very similar experimental design ^(18)^.

Demonstration of delayed fracture repair in OVX C3H/HeJ mice is a strength of this study as it replicates the up to 50% of fragility fractures do not heal in expected time frames ^(39-42)^. There has been variable success in replication of post-menopausal osteoporosis delayed fracture healing in rat models ^(60,61)^, with OVX having to be combined with calcium deficiency to achieve robust presentation of this complication ^(62)^. Whereas OVX-associated delayed healing in mouse fracture studies have been limited by experimental approach. Specific confounders include fracture and OVX surgeries being performed at a similar time rather than allowing osteopenia to develop prior to fracture ^(63,64)^; pre-fracture OVX being performed in young growing mice ^(65)^; or use of a partial bone defect rather than fracture model ^(66)^. In several studies with an experimental approach appropriate for modelling post-menopausal fragility fracture, the fracture repair outcomes were mixed and so did not conclusively support delayed healing ^(42,67)^. Both choice of mouse strain and reproducibility of fracture fixation were likely advantages that facilitated robust detection of delayed fracture healing herein. Overall, OVX in C3H/HeJ adult mice replicates major features of human post-menopausal osteoporosis, validating its use for exploration of novel pathophysiology of this disease.

Given the critical role of osteomacs in supporting osteoblast-mediated bone anabolism and that inappropriately low or reduced osteoblast function is a hallmark of osteoporosis, the OVX-associated increase in osteomacs observed herein was an unexpected outcome. An obvious interpretation is osteomacs are increased to supply the estrogen-deficiency-induced elevated demand for osteoclast precursors. While *in vitro* studies support this interpretation ^(27,68)^, it is challenged by *in vivo* studies indicating that macrophages do not function efficiently in this capacity ^(69,70)^. Notably, in mice treated with an exogenous supra-physiologic RANKL, osteomacs remained TRAP-negative and continued to be a prominent cell within the endosteal micro-environment despite substantially elevated osteoclastogenesis ^(71)^. Therefore, osteomacs may have distinct contributions to osteoporosis pathophysiology beyond serving as an osteoclast precursor.

The increase in osteomacs could be a compensatory mechanism to correct the OVX-associated reduction in osteoblast number and/or function. However, in situ assessment highlighted osteomac direct association with osteoclasts, positioned adjacent to the basolateral membrane that contains the FSD. Co-localisation of osteomacs and osteoclasts has been reported previously ^(29,71,72)^ without interrogation of the purpose of this anatomical distribution. Notably, CD169^+^ osteomac/macrophage depletion increased TRAP^+^ subcellular deposits in bone marrow interstitium, as shown herein, and elevated serum TRAP, as per previously. Both of these indicators increased in the absence of any change in osteoclast number, area, TRAP staining intensity or osteoclast function ^(30)^. This suggests that in the absence of osteomac/macrophage, bone resorption by-products accumulate in bone marrow and diffuse into the circulation. During osteoclastic bone resorption, degraded bone matrix and associate components are endocytosed and transported to the basolateral membrane and expelled into the extracellular fluid through the FSD ^(53)^. Intriguingly, osteoclast-adjacent osteomacs frequently contained TRAP^+^ subcellular vesicles. TRAP has been shown to be abundant within intracellular endocytic vesicles in osteoclasts actively resorbing bone ^(73)^ supporting that TRAP would be present in exocytosed post-resorption vessicles. In combination with their physical location, this suggests that osteomacs phagocytose bone resorption by-products being expelled from osteoclasts. Further evidence supporting osteomac-mediated ‘clean-up’ of bone resorption by-products was demonstrated using *in vitro* cultures in which macrophages contained devitalised bone fragments only when co-cultured with osteoclasts.

While it is possible that osteomacs could be endogenously expressly TRAP, as some macrophage populations have been shown to express acp5 mRNA ^(74)^, the collective evidence presented herein supports that osteomacs are functioning to eliminate, or at least sequester, post-bone resorption by-products. Osteomac-mediated post-resorption clean-up, combined with the demonstrated increase in osteomac number, provides a likely explanation for why elevated serum TRAP, as a maker of increased bone resorption, was not observed in the C3H/HeJ OVX model. Serum TRAP is an inconsistent marker of bone catabolism in murine OVX models ^(75-77)^. It would be of interest to assess whether OVX-induced osteomac increased frequency is blunted in models in which elevated serum TRAP has been detected. Given post-bone resorption by-products potentially contain bio-active molecules able to influence skeletal or hematopoietic homeostasis, osteomac supporting osteoclast-mediating bone resorption in this manner could restrict local and/systemic unintended impacts of bone resorption. Presence of TRAP+ intracellular vesicles within osteomacs was reported herein in growing rat bone and previously in rapidly remodelling periosteal callus associated with bone injury ^(29)^. Thus, it is likely the osteomacs function in this capacity not only during pathology-associated elevated bone resorption, but also during normal bone catabolic modelling and remodelling associated with growth and healing.

Given the collective evidence implicates osteomacs in supporting both bone anabolism and catabolism, we examined the skeletal impacts of loss of the macrophage-restricted molecule, CD169 (Siglec-1). We observed an age-associated low bone mass phenotype in male CD169-KO mice. While CD169 has not been directly implicated in contributing to bone biology or skeletal homeostasis, it does contribute to macrophage phagocytic function ^(78)^ and cell-cell adhesion ^(79)^. Therefore, there is the potential for CD169 to participate in osteomac support of both osteoclasts and osteoblasts ^(34)^. Further interrogation of CD169 in mediating osteomac phagocytosis of bone resorption by-products, phagocytosis of apoptotic osteoblasts and/or direct interaction with osteoclasts and/or osteoblasts is needed. The cell dynamics associated with CD169 deficiency showed reduced osteomac frequency at 8 weeks while decreased osteoblasts and increased osteoclasts were detected at 16 weeks. The total myeloid population is minimally affected in the CD169-KO mice, supporting that the bone impacts are not due to limited OC precursor availability. However, the reduction in osteomac number supports that this contributed, at least in part, to fractional bone loss. A more detailed time course study would be needed to dissect the disconnect in cellular dynamics underpinning this phenotype and whether loss of CD169 compromised attainment of peak bone mass or contributed to development of osteopenia.

In conclusion, the comprehensive validation of OVX in adult C3H/HeJ mice will be valuable in guiding future studies exploring novel osteoporosis pathological mechanisms and interventional studies, including those targeted at correcting delayed fracture healing. Furthermore, we exposed a novel role for osteomacs in osteoclast-mediated bone resorption and provide the first evidence of their involvement in osteoporosis bone pathology. We also demonstrated for the first time that the loss of expression of a single macrophage-specific molecule is sufficient to cause a low bone mass phenotype. Overall, this study extends on the accumulating evidence that osteomacs, through influence of both catabolic and anabolic bone mechanisms, are important contributors in attainment and maintenance of peak bone mass.

## Authors contribution

LB: experimental design, experimental execution, data collection, data analysis, figure generation. SMM and LJR: funding acquisition, experimental design, experimental execution. AW; SK: assisted in experimental execution. LWHS, KW, CS: assisted in data analysis. MB and VG: assisted in data collection. NJP and PYN: funding acquisition, experimental design, experimental execution, data collection, data analysis. ARP: funding acquisition, experimental design, experimental execution, data analysis and led and coordinated research. LB and ARP wrote the manuscript and all other authors reviewed, edited and/or approved manuscript.

## Acknowledgements

This work was supported by Mater Foundation and a National Health and Medical Research council Project Grant (APP1143802), The University of Queensland Postgraduate Scholarship and Mater Research Frank Clair Scholarship (LB), an Australian Research Council Future Fellowship (ARP, FT150100335), an Arthritis Australia and HJ & GJ Mackenzie Grant (NJP and PYN), and a Faculty of Health and Medical Sciences Research Grant Scheme (SE Ohman Medical Research Fund; NJP and PYN). This work was carried out at the Translational Research Institute (TRI) which is supported by a grant from the Australian Government. The Translational Research Institute (TRI) Microscopy, Histology, Preclinical Imaging and Flow Cytometry Core Facilities contributed technical expertise and the TRI Biological Research Facility contributed to animal husbandry and monitoring. The authors acknowledge the facilities, and the scientific and technical assistance of Microscopy Australia at the Centre for Microscopy, Characterisation & Analysis, The University of Western Australia, a facility funded by the University, State and Commonwealth Government. The rat tissues were generously provided by Professor David Hume and Dr Katharine Irvine. Professor Martin Wullschleger and Associate Professor Ben Ahern provided surgical expertise related to MouseScrew fracture generation.

## Notes

### Competing Interest Statement

The authors have declared no competing interest.

