## Supplemental Figures for "Osteomacs support osteoclast-mediated resorption and contribute to bone pathology in a postmenopausal osteoporosis mouse model"

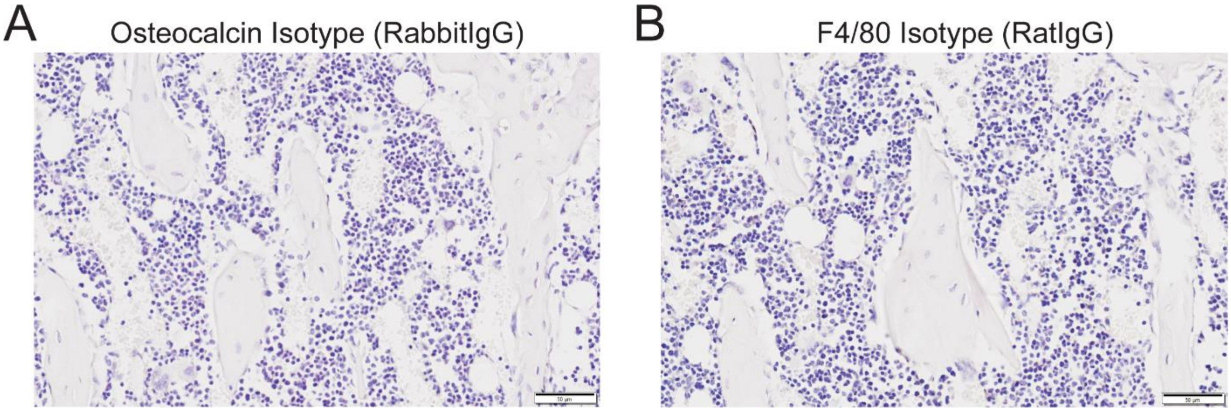


**Supplemental Figure 1. IHC isotype antibody control staining.** Representative images of isotype antibody control staining to demonstrate specificity of staining for osteocalcin (**A**) and F4/80 (**B**). Images show metaphyseal trabecular bone region of proximal tibia. Original magnification: 400X. Scale bar: 50 µm.

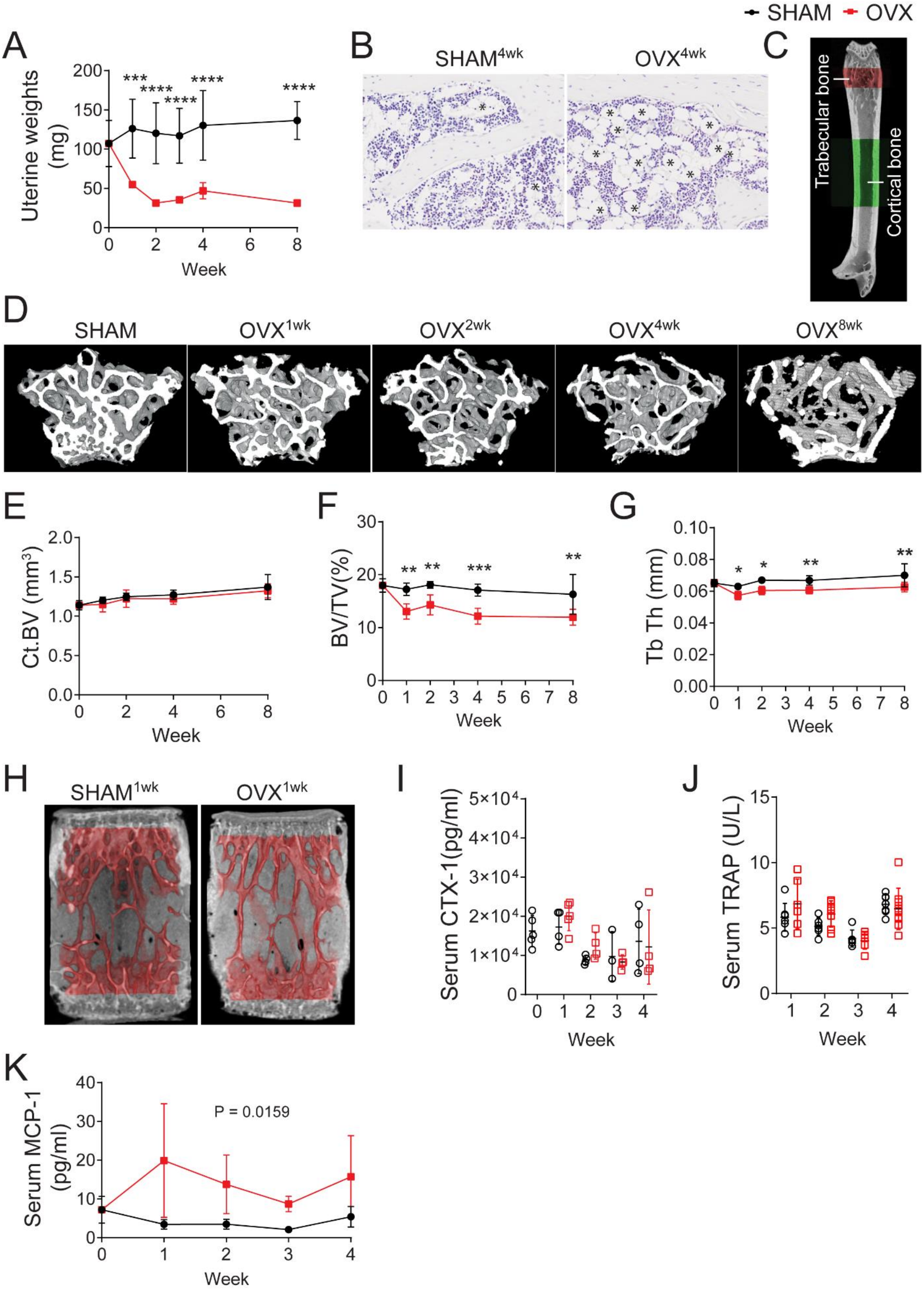
SHAM
OVX
A
Uterine weights (mg)
Week
B
SHAM4wk
OVX4wk
C
Trabecular bone
Cortical bone
D
SHAM
OVX1wk
OVX2wk
OVX4wk
OVX8wk
E
Ct.BV (mm3)
Week
F
BV/TV(%)
Week
G
Tb Th (mm)
Week
H
SHAM1wk
OVX1wk
I
Serum CTX-1(pg/ml)
Week
J
Serum TRAP (U/L)
Week
K
Serum MCP-1 (pg/ml)
P = 0.0159
Week

**Supplemental Figure 2. OVX in C3H/HeJ recapitulates features of human osteoporosis. A:** Uterine weight loss post-OVX across an 8-week time course. **B:** Representative histological images from hematoxylin-stained sagittal sections of the tibia showing adipocytes (asterisks) based on morphological appearance in the bone marrow at 4 weeks post-SHAM or OVX surgery. **C:** Micro-CT 3D reconstruction of femur showing regions of interest analysed in the metaphyseal trabecular (red) and diaphyseal cortical (green) bone. **D:** Representative micro-CT images of femoral trabecular bone exemplifying bone loss progression over 8 weeks post-surgery. **E:** Micro-CT analysis of cortical bone volume (Ct.BV). **F-G:** Micro-CT analysis of vertebral trabecular bone volume per total volume (BV/TV) (**F**) and trabecular thickness (Tb.Th) (**G**). **H:** Representative micro-CT 3D reconstructions of vertebrae from SHAM- or OVX-operated animal showing the region analysed masked in red. **I-J:** Serum CTX-1 and TRAP levels after SHAM or OVX operation. **K:** Serum levels of monocyte chemokine attractant protein 1 (MCP-1) in SHAM- or OVX-operated mice at different weeks post-OVX. Statistical significance was determined using two-way ANOVA with Tukey's post-test. ****, $p < 0.0001$; ***, $p < 0.001$; **, $p < 0.01$; *, $p < 0.05$. P-value in **K** shows significance due to surgery (OVX). Error bars represent standard deviation. Original magnification: 200X.
